# Natural strongyle infection reduces relative abundance of inflammation-inducing *Prevotella* in wild primates

**DOI:** 10.1101/2022.06.02.494558

**Authors:** Carrie A. Cizauskas, Alex D. Washburne, Joseph E. Knelman, Christina B. Hansen, Antony Mwangi Nderitu, Peter Lokwamo Esinyon, Andrew P. Dobson, Andrea L. Graham

## Abstract

Microbes living within the mammalian gastrointestinal tract affect the metabolization and extraction of dietary nutrients, immune function, colonization by pathogens, and risk of autoimmune disease. While most microbiome studies focus on sequences of the 16S gene shared by Bacteria and Archaea, these are not the only regular inhabitants of mammalian guts. Macroparasites such as helminths are nearly ubiquitous in wildlife, and a quarter of the world’s human population harbors helminths; these worms affect host physiology as they compete with microbiota over host resources while also affecting host immunity, and changing the host microbiome. Little is understood about how helminths interact with microbiomes to affect host disease states, and few studies have examined these interactions in natural systems in genetically diverse hosts experiencing coinfections and other stressors.

We surveyed the microbiomes and helminth parasites of wild primates and found strong associations between helminths and microbes in the bacterial microbiome. Notably, we find that the presence of a strongyle we hypothesize to be hookworm is correlated strongly with decreased relative abundance of *Prevotella* species, a lineage associated with inflammatory bowel disease humans. This observed decline in *Prevotella* relative abundance, a genus implicated in several host autoimmune and inflammatory disorders, motivates future research on whether the mixed results of helminthic therapy (*i*.*e*., “infecting” patients with gastrointestinal nematodes to treat various diseases) stem from the mixed causes of inflammation, and whether inflammation specifically correlated with *Prevotella*-driven dysbiosis can be mediated through mechanisms mimicking how hookworms and other nematodes behave in the gastrointestinal ecosystem of their hosts. Our findings lend ground-truthed support to previous lab-based studies and limited/restricted human trials showing potential benefits, via microbial modulation, of nematode therapy in treating inflammatory bowel disease. Our study adds statistical weight to a link between helminths and a specific lineage of microbes associated with inflammation.

## Introduction

While much is being learned about how gastrointestinal (GI) microbes affect host health and physiology, relatively little is known about how ‘macrobes’ such as helminths interact with the microbiome to affect hosts. Microbes help extract and metabolize nutrients from host diets[1], develop and regulate immune function[2], and protect against entero- and other pathogens[3]. Research suggests that host gut ecosystems are similar to traditional ecosystems: they are more robust to perturbation when they harbor higher (microbial) diversity[4]; lower levels of perturbation are synonymous with the good health of their “owners”. Microbiome dysbioses have been correlated with, and even implicated in, the development and exacerbation of metabolic disorders: obesity, malnutrition, diabetes, some cancers, some neurological disorders, and several chronic inflammatory diseases (*e*.*g*. Crohn’s disease, ulcerative colitis, celiac disease)[5]. Microbes found by sequencing the 16S gene, however, are only a subset of organisms common to the mammalian gut biome.

While GI helminth and protozoal parasites have co-evolved over millennia with humans and other mammals[6], and while they are nearly ubiquitous as co-infections in wildlife populations[7], only one-third of the global human population still harbors helminths[8,9]. The relatively recent decrease in human-macroparasite interactions has been hypothesized to adversely affect GI microbial diversity, with downstream effects on host health and immune dysregulation[10].

Relatively few studies have examined macroparasite-microbiome interactions in mammalian hosts, most of which have been done with laboratory animals harboring single species helminth infections[11–17]. However, these studies have found a clear influence of helminths on mammalian gut microbiomes. To our knowledge, only two previous studies have examined parasite-microbiome interactions in genetically-diverse mammalian hosts (*i*.*e*., not inbred laboratory animals) in natural systems[18],[19]. In this paper, we examine *in situ* associations between microbiomes across two wild primates species and 10 genera/species of helminths, to identify whether changes in microbial lineages predictably occur in association with helminth infections.

Non-human primates (NHP) are a model system for observational studies of the effects of diet and parasites on host microbiomes and their interaction effects on human health. Wild NHP are particularly important to study, as they provide close genetic proxies to humans dealing with coinfections, dietary changes, hormonal oscillations, social pressures, and other natural stressors that fluctuate seasonally. Our study species, *Papio anubis* (olive baboons) and *Chlorocebus pygerythrus* (vervet monkeys), are well studied both in laboratory and natural settings. Because these species remain in relatively stable groups with predictable ranges and sleeping sites, we were able to control for genetic and behavioral variation within groups and obtain large sample sizes. We evaluated 366 fecal samples from baboons and 327 fecal samples from vervets (Table 1) collected over three seasons in one year for parasite genera/species presence, infection intensity, and richness. We identified five clades of parasites partitioned into 13 presumptive (identifiably different) genera/species, 10 of which were helminths.

**Table 1.** Total fecal samples collected for each study host species over three sampling seasons.

| Troop Identity | Sp. | Season | Number of Fecal Samples Collected | Troop Size | % Troop Sampled for Parasites | Number of Fecal Samples Sequenced | % of Fecal Samples Sequenced | % of Troop Sequenced for Microbiome Analysis |
| --- | --- | --- | --- | --- | --- | --- | --- | --- |
| Cliffs | PA | Jan | 27 | 70 | 38.6 | 7 | 25.9 | 10.0 |
| Cliffs | PA | Apr-May | 49 | 65 | 75.4 | 15 | 30.6 | 23.1 |
| Cliffs | PA | Nov-Dec | 35 | 62 | 56.5 | 28 | 80.0 | 45.2 |
| Meg's | PA | Jan | 2 | 42 | 4.8 | 0 | 0.0 | 0.0 |
| Meg's | PA | Apr-May | 48 | 46 | 100.0+ | 25 | 52.1 | 54.3 |
| Meg's | PA | Nov-Dec | 25 | 46 | 54.3 | 16 | 64.0 | 34.8 |
| Small | PA | Jan | 0 | 30 | NA | 0 | NA | NA |
| Small | PA | Apr-May | 22 | 30 | 73.3 | 7 | 31.8 | 23.3 |
| Small | PA | Nov-Dec | 24 | 30 | 80.0 | 14 | 58.3 | 46.7 |
| Limpy's | PA | Jan | 44 | 55 | 80.0 | 15 | 34.1 | 27.3 |
| Limpy's | PA | Apr-May | 0 | 55 | NA | 0 | NA | NA |
| Limpy's | PA | Nov-Dec | 0 | 55 | NA | 0 | NA | NA |
| Shorttail's | PA | Jan | 0 | 45 | NA | 0 | NA | NA |
| Shorttail's | PA | Apr-May | 0 | 45 | NA | 0 | NA | NA |
| Shorttail's | PA | Nov-Dec | 31 | 45 | 68.9 | 12 | 38.7 | 26.7 |
| Shorttail's + Limpy's together | PA | Jan | 11 | 100 | 11.0 | 4 | 36.4 | 36.4 |
| Shorttail's + Limpy's together | PA | Apr-May | 0 | 100 | NA | 0 | NA | NA |
| Shorttail's + Limpy's together | PA | Nov-Dec | 0 | 100 | NA | 0 | NA | NA |
| Naibor<br>Cattle Dip | PA | Nov-Dec | 36 | ~80 | ~45 | 14 | 38.9 | 17.5 |
| Naibor<br>Garden | PA | Nov-Dec | 12 | ~30 | ~40.0 | 9 | 75.0 | 30.0 |
| Fig | CP | Jan | 31 | 23 | 100.0+ | 9 | 29.0 | 39.1 |
| Fig | CP | Apr-May | 3 | 23 | 13.0 | 3 | 100.0 | 13.0 |
| Fig | CP | Nov-Dec | 17 | 23 | 73.9 | 10 | 58.8 | 43.5 |
| Hippo<br>Pool | CP | Jan | 21 | 32 | 65.6 | 12 | 57.1 | 37.5 |
| Hippo<br>Pool | CP | Apr-May | 28 | 32 | 87.5 | 9 | 32.1 | 28.1 |
| Hippo<br>Pool | CP | Nov-Dec | 26 | 32 | 81.3 | 15 | 46.9 | 46.9 |
| Bridge | CP | Jan | 3 | 14 | 21.4 | 1 | 33.3 | 7.1 |
| Bridge | CP | Apr-May | 40 | 14 | 100.0+ | 13 | 37.5 | 92.9 |
| Bridge | CP | Nov-Dec | 15 | 14 | 100.0+ | 10 | 66.7 | 71.4 |
| Ranch | CP | Jan | 20 | 30 | 66.7 | 10 | 50.0 | 33.3 |
| Ranch | CP | Apr-May | 14 | 30 | 46.7 | 4 | 28.6 | 13.3 |
| Ranch | CP | Nov-Dec | 16 | 30 | 53.3 | 11 | 68.8 | 36.7 |
| Kudu | CP | Jan | 0 | 28 | NA | 0 | NA | NA |
| Kudu | CP | Apr-May | 70 | 28 | 100.0+ | 17 | 24.3 | 60.7 |
| Kudu | CP | Nov-Dec | 15 | 28 | 53.6 | 12 | 80.0 | 42.9 |
| MRC | CP | Jan | 0 | 7 | NA | 0 | NA | NA |
| MRC | CP | Apr-May | 7 | 7 | 100.0 | 4 | 57.1 | 57.1 |
| MRC | CP | Nov-Dec | 1 | 4 | 25.0 | 1 | 100.0 | 25.0 |
| Total | PA | Jan | 84 | 242<sup>1</sup> | 34.7<sup>2</sup> | 26 | 40.0 | 10.7 |
| Total | PA | Apr-May | 119; 117* | 241 | 49.4; 48.5 | 47 | 39.5 | 19.5 |
| Total | PA | Nov-Dec | 163 (115 MRC only) | 348 (238 MRC only) | 46.8 (48.3 MRC only) | 93 (70 MRC only) | 57.1 (60.9 MRC only) | 26.7 (29.4 MRC only) |
| Total | CP | Jan | 75; 67* | 134 | 56.0; 50.0 | 32 | 42.7 | 23.9 |
| Total | CP | Apr-May | 162; 94* | 134 | 100.0+; 70.1 | 52 | 32.1 | 38.8 |
| Total | CP | Nov-Dec | 90; 89* | 131 | 68.7; 67.3 | 59 | 65.6 | 45.0 |

Sp. refers to species, where PA is *Papio anubis* (olive baboon) and CP is *Chlorocebus pygerythrus* (vervet monkey).

Season refers to sampling season, where Jan. refers to the pilot season in January, 2014, Apr-May refers to the five week sampling season in April-May, 2014, and Nov-Dec refers to the four week sampling season in November-December, 2014.

Number of Fecal Samples Collected, Number of Fecal Samples Sequenced, and % of Fecal Samples Sequenced is ultimately sometimes lower than one third of the troop at each time point, as this number is based on the final number of samples sequenced after quality control methods were implemented, and not on the original goal of sequencing at least approximately one third of the samples collected from each troop at each sampling time point.

100.0+% refers to groups for which more samples were collected during that sampling period than there were members of the troop, indicating some resampling of individuals.

^1^Sizes of the total populations of baboons (for all five troops) or vervets (for all six troops) at each sampling period. We used only the numbers for Limpy’s & Shorttail’s Together for the population counts for those troops (100 for the combined group for each of the three seasons). *Total samples with two numbers list first total samples for that species at that sampling period, followed by the total sample numbers assuming the most minimal amount of individual repeat sampling (e.g. although 40 samples were collected from the Bridge Vervet group in April-May, there were only 14 animals in this troop, so only 14 samples were counted in this second tally). ^2^Percent of total populations for each species sampled at each sampling period. For entries with two numbers, the first number is percent of population sampled using total sample numbers, while the second number is percent of population sampled using the numbers for most minimal amount of individual repeat sampling.

We extracted DNA from 166 baboon fecal samples and 143 vervet fecal samples (see Methods), and performed Illumina MiSeq sequencing with 16S libraries using primers for the 16S rRNA V4 region. Phylofactorization[20,21] of the microbial community dataset identified major axes of variation in the microbial community attributable to host species, diet, season, and helminth genera/species presence/absence.

## Results and Discussion

The dominant phylogenetic factors in the microbiome clade community composition distinguished between host species and corresponded to phylogenetic diet patterns observed in humans in the American Gut Project[20]. Controlling for effects of diet and season, helminths were associated with differences in the relative abundances of many microbial clades even when controlling for multiple hypothesis tests inherent in this phylogenetic analysis. *Trichuris* and hypothesized hookworms (Strongyle #2; hereafter referred to as “hookworms” to indicate hypothesized status) generally showed opposite effects on clade relative abundances; for example, “hookworm” infection was associated with decreases in relative abundance of *Prevotella* and *Bacilli*, whereas *Trichuris* infection resulted in no decrease in these microbes compared to “controls” (animals without *Trichuris*) (Figs 1 and 2). Across phylogenetic factors, we found an unexpected pattern: “hookworms” and tapeworms tended to have opposite effects on microbial lineages’ relative abundances (Fig 2).

**Figure 1.**
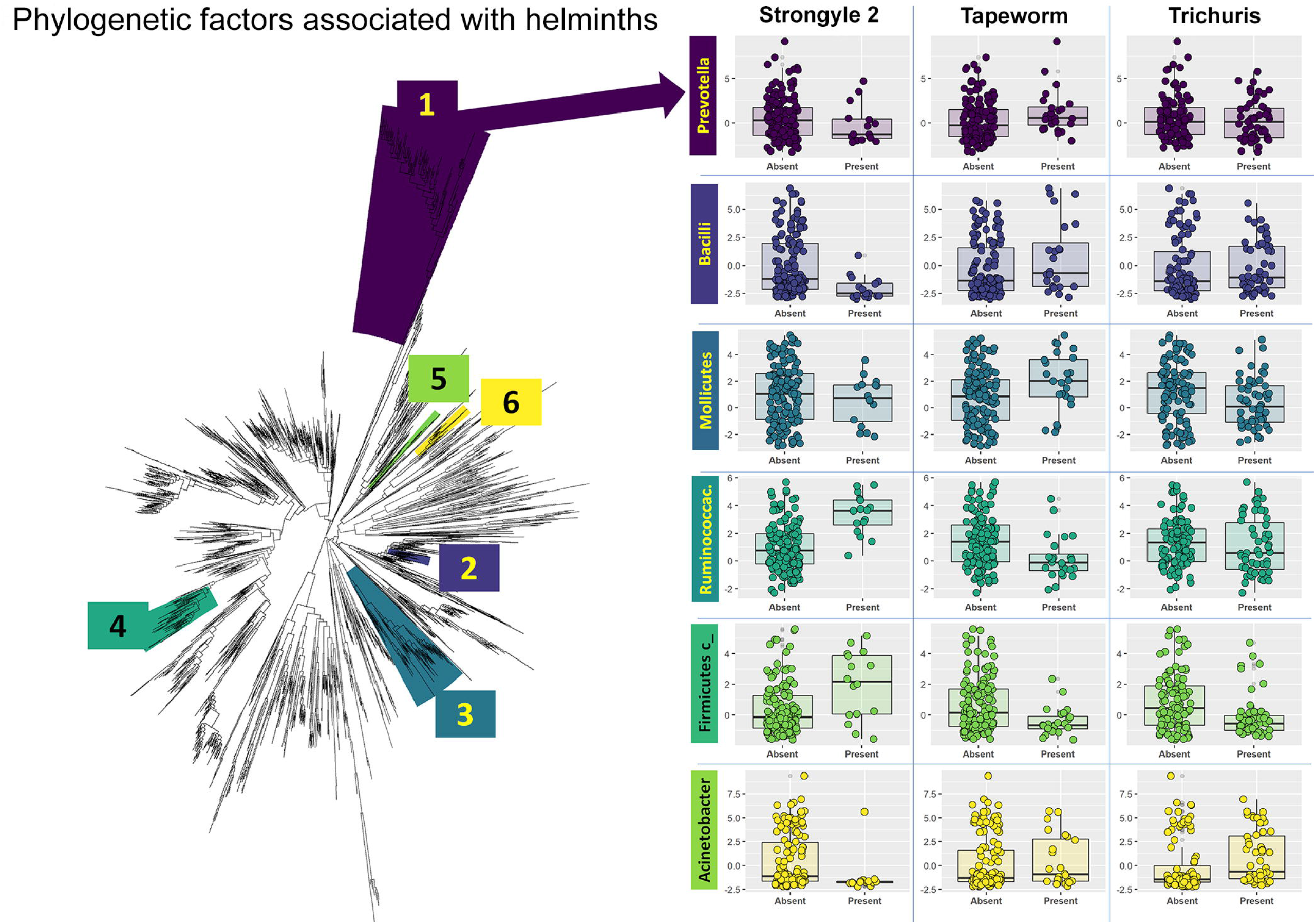
Phylogenetic factors of helminth infections. Helminth infections affect the relative abundances of many large clades of microbes. Fifty phylogenetic factors (lineages) yielded six large (≥ 5 OTU) clades of microbes with differential relative abundances associated with helminth infections. Boxplots of the clades’ isometric log-ratios (ILR) abundances versus the presence and absence of three main helminths are plotted, with colors of boxplots corresponding to the clades highlighted on the phylogeny. The first major clade impacted by helminths was a clade of 100 OTUs of the genus *Prevotella* found to decrease in the guts of primates infected with “hookworms” (Strongyle 2). Note that ILR abundances are a transform for relative abundances; we use ILR rather than logic transformation for microbiome data analysis because of the geometric nature of microbial variation (the tendency of the variance to increase like the square of the mean, the tendency of attrition processes like variable rates of lysis to multiplicatively decrease abundances and PCR amplification biases to multiplicatively increase abundances). Phylofactorization constructs a complete ILR basis orthogonal to **1**, with each element corresponding to a lineage in the phylogeny.

**Figure 2.**
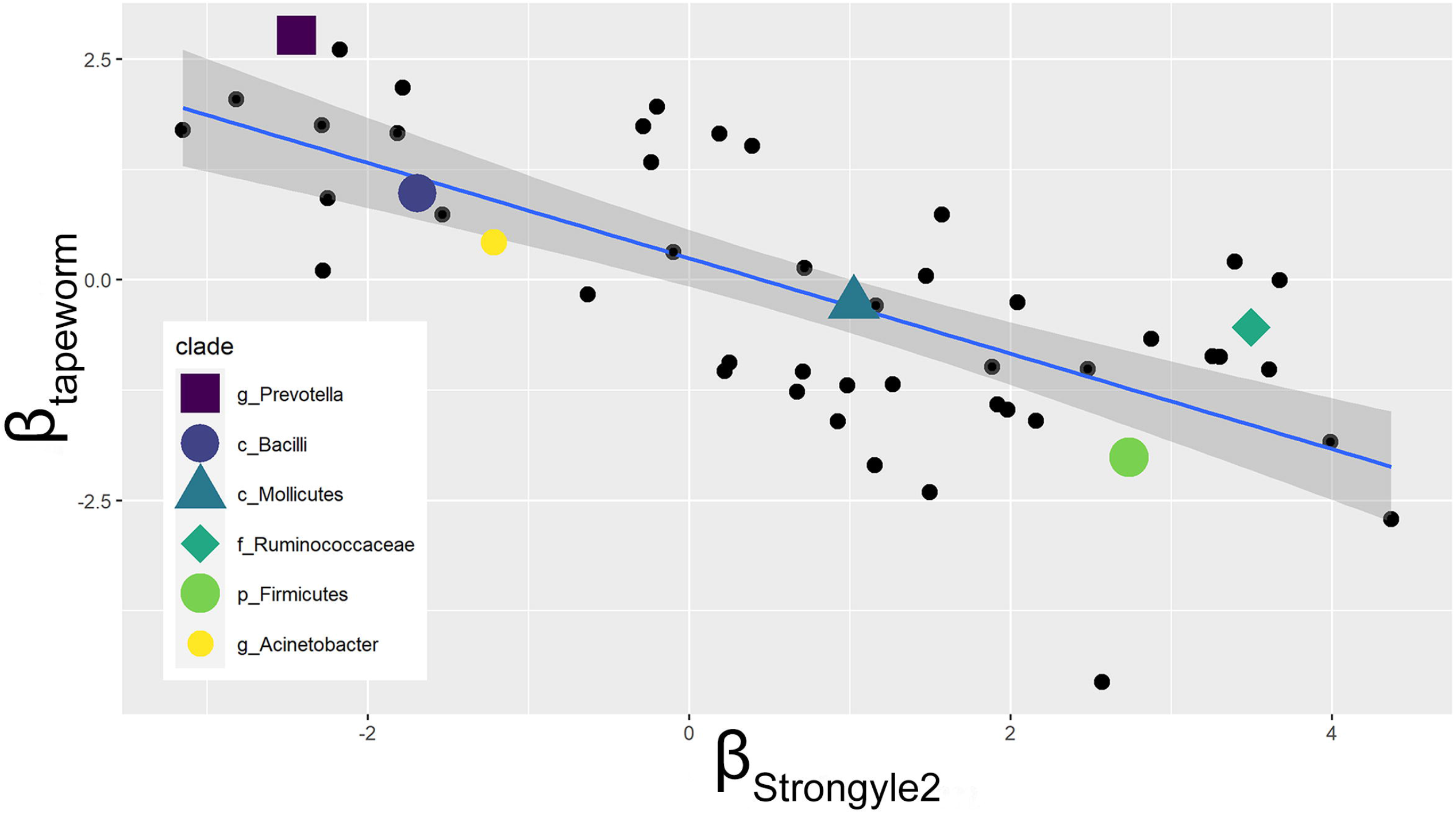
Opposing effects of Strongyle #2 (presumptive hookworms) and tapeworms on primate microbiomes. Across all 50 phylogenetic factors of bacteria associated with helminth infections (controlling for primate species and time of year), lineages whose relative abundances were positively associated with hypothesized hookworms (Strongyle 2) tended to have relative abundances negatively associated with tapeworms, and vice-versa. Shown here is the negative relationship between the relative log-ratio of geometric mean abundances of these taxa within the group *vs*. all other taxa.

Our finding that helminths interact with microbiome composition is in concert with previous studies, while being the first to examine these interactions in higher primates. Previous investigations have found varying interaction effects: increased bacterial diversity associated with presence of specific parasites[11,13,18]; parasites associated with reduced microbial diversity or dysbiosis[14,16,22]; decreased bacterial abundance with the presence of specific parasites[12,18]; and increased microbial diversity or widespread changes in the presence of parasite infection[18,19,23,24]. We found only one study that reported no changes in microbiome composition in conjunction with helminth infection[25]. While mock communities may refine our findings by eliminating potential contaminants, our use of methods control for compositional effects, as well as differences across species and season of sample collection. Thus, our findings of significant changes in abundant microbial lineages that are well known to inhabit the mammalian gut suggest these findings reflect real changes in community composition associated with helminthic cohabitants of the gut.

Intriguingly, we found a clade of 100 OTUs of genus *Prevotella* decreased in primates infected with presumptive hookworms (Fig 1). *Prevotella*, gram-negative bacteria in phylum Bacteroidetes, are typically enriched in hosts consuming high fiber diets, as these bacteria help digest complex plant polysaccharides[26]. However, some *Prevotella* are known to cause opportunistic infections and provoke inflammatory responses compared to strictly commensal organisms[27]. Increased relative abundance of various *Prevotella* species in human guts has been associated with localized and systemic inflammatory and autoimmune diseases, including insulin resistance and obesity[28], HIV-associated GI and systemic inflammation[29], ulcerative colitis and Crohn’s disease[30], systemic sclerosis[31], ankylosing spondylitis[32], periodontitis[27], colon cancer[33], and rheumatoid arthritis (RA)[34]. Scher *et al*.[34] found that significant enrichment of *Prevotella copri* was strongly correlated with noticeable RA symptoms in 75% of patients with new-onset, untreated RA, and with a loss of beneficial microbes in these patients; these findings were corroborated by others[35]. Mice that were first treated with antibiotics, then given *P. copri* orally, rapidly developed a *P. copri*-dominated gut environment and experienced significantly more severe colitis, weight loss, gut epithelial damage, and inflammatory cytokine production when compared to controls exposed to a colitis-inducing substance[34].

A few previous studies have demonstrated reduced Prevotellaceae abundances in host guts with concurrent infections of whipworm[15], strongyles[36], and amoebozoans[8]. Our study is the first to correlate decreased *Prevotella* relative abundance with hypothesized hookworm infection. Increased *Prevotella* abundance may lead to pathology, as these bacteria can degrade and decrease mucin production[15]; through these effects on this gut wall protector, or through direct actions, *Prevotella* strongly induces inflammatory mediators such as caspase-1, NF-kappaB, and Th17-related cytokines[15,35], leading to both mucosal and systemic inflammation (Larsen 2017).

The mechanisms of interaction between hookworms (and other nematodes) and *Prevotella* are still being examined. Helminths may alter microbiota through competition for nutrients[15,37], or through excretory/secretory products (ESP) that alter the intestinal microbial habitat and/or protect the gut from inflammation[37]. Various helminths have been long known to secrete and/or excrete short chain fatty acids (SCFA)[38,39] and small molecules[39–41]. Many of these small molecules appear to have anti-inflammatory and other pharmacological properties[39–41]. SCFA have long been known to increase colonocyte proliferation, aid colonocyte function, promote adaptive gut responses to intestinal disturbances and injury, increase gut blood flow and motility, and decrease GI auto-immune pathology by inducing anti-inflammatory cytokine secretion and enhancing local regulatory T cell numbers and function[42–44]. Gut bacteria can also produce SCFA when they degrade starch and fiber in dietary foods[43]. Perhaps most interestingly, recent research indicates that gut helminths and microbes may interact to increase SCFA production: Zaiss *et al*.[44] found that intestinal nematodes in mice caused GI bacterial microbiota to increase production of SFCA. These “activated” microbes then continued to produce increased levels of SCFA even in the absence of helminths; in both treatments (helminths present; mice treated with “activated” microbes alone), the SCFA interacted with regulatory T cell receptors to ultimately decrease allergic symptoms in asthmatic mice.

Helminths have also been found to polarize the immune system toward a Th2-type response, suppress inflammatory Th1- and Th17-type responses, and reduce immunopathology[5,36]. Helminths may do this by promoting regulatory and tolerogenic immune crosstalk through their SCFA production and/or induction of SCFA production by microbes (see above)[44]. One study found that mice infected with hookworm *Nippostrongylus brasiliensis* had significantly lower transcript levels of all tested IL-17-associated genes in ileal tissues[36]. Others have found that this parasite stimulates mast cells through IL-4 induction, thereby increasing innate responses to bacteria[45], and presence of these parasites has been correlated with a more than 2000-fold decrease of segmented filamentous bacteria known to promote Th17 differentiation and induce inflammation-associated genes.[36] Helminth-driven Th2-skewing can decrease immune responses to bacterial antigens[45], and helminth Th2-induction may decrease bacterial translocation across gut epithelium, leading to decreased systemic inflammation[46]. Interestingly, studies of humans in rural Africa and Asia have found high relative abundances of *Prevotella* in host guts, likely due to consumption of diets rich in starch, fiber, and plant polysaccharides[30], yet these populations have very low prevalences of autoimmune disorders. We hypothesize that a higher prevalence of gut macroparasites in these groups may mitigate the potential inflammatory effects of *Prevotella*. Scher *et al*.[34] hypothesized that *P. copri* may thrive in a pro-inflammatory environment and may exacerbate inflammation for its own benefit, a *milieu* that may be controlled by the presence of helminths.

Helminth therapy for inflammatory/autoimmune diseases has been explored with promising results in humans with celiac disease[47], mice with IBD[48], and NHP with idiopathic chronic diarrhea[12]. Hookworms may be particularly good at decreasing autoimmune-based inflammation; experimental infections with small numbers of *Necator americanus*, the most widely distributed human hookworm, have been both well-tolerated by hosts and helpful in treating celiac disease, asthma, and other inflammatory ailments.[5] Helminths may also affect host immunity and inflammation through direct effects on the microbiome[49]. They may also indirectly improve host health by supporting increased gut microbial richness[4]; for example, there is evidence that *N. americanus* infection in particular may lead to increased gut microbiome diversity[23]. Helminth therapy may therefore be useful not only for its anti-inflammatory effects in the face of autoimmune disease, but could be envisioned as a method for regulating other aspects of host health.

Scientists have recently suggested that a diverse “multibiome,” including cohabitating bacteria, viruses, fungi, and macroparasites, may be necessary components of a healthy mammalian gut environment[50]. Some have hypothesized that the increase in autoimmune and chronic inflammatory disorders in industrialized countries may be directly due to a lack of helminths in this multibiome[9], as helminths can be both anti-inflammatory and increase immune gene diversity in their human hosts[51]. Others have suggested that decreased prevalence of gut helminths and protozoa in industrialized countries may be partly responsible for declines in GI bacterial diversity in these populations, which may lead both directly and indirectly to host health issues[10]. This bacterial diversity may be restored therapeutically by helminth treatment[12,52]. Our findings suggest that helminth therapy may be a nuanced tool, with particular taxa of helminths capable of mediating autoimmune or microbial-induced pathologies. By identifying precisely which microbial lineages show differential relative abundances under the presence of helminth infections in wild primates (Fig 1), and showing the opposite effects different helminths sometimes have on bacterial relative abundances (Fig 2), we illuminate the nuanced niche of helminths in the community structure of wild primate multibiomes. Our study leads the way for further research on the interactions of specific species of gastrointestinal helminths and the gut microbiome in natural populations and populations with parasitic coinfections.

## Methods

### Study area and species

We conducted field research at the Mpala Research Center (MRC) and the associated Mpala Ranch, which are located on 190km^2^ of land within the Laikipia district of central Kenya (37°53′ E, 0°17′ N). Field research was approved by the Mpala Research Center; the Kenya Wildlife Service (KWS/BRM/5001); the Kenya National Commission for Science, Technology and Innovation (NACOSTI/P/14/9392/1023); the Kenya Veterinary Board (KVB/GEN/Vol.II/328); and the Kenya National Environment Management Authority (0045). Sample exportation and importation were approved by the U.S. Centers for Disease Control and Prevention (2015-03-049); the Kenya Wildlife Service; and Princeton University. We expanded the study in the final season to include two sites near the Nanyuki River in Naibor (approximately 16km from MRC). The study areas consist of semi-arid savanna and riverine woodland with rolling hills and granite outcroppings. Mean annual rainfall 1998-2015 in MRC was 643.3mm (+/- 194.4mm); total rainfall during the study year was 443.2mm. The area historically experienced delineated dry, long rainy, and short rainy seasons corresponding to our sampling seasons[53], though these seasons have been less clear cut in recent years.

Study subjects were seven known groups of olive baboons (*Papio anubis*; two of these groups, Limpy’s and Shorttail’s, would sometimes combine into an eighth “supergroup” that could only be found in this form during certain seasons – see Table 1) and six known groups of vervet monkeys *(Chlorocebus pygerythrus*). Primate sampling was approved by the Princeton University IACUC board (Number 1972A-14). Sampling was conducted completely non-invasively. Two of the baboon groups were added in the final sampling season as opportunity arose, and as such were not sampled until the Nov/Dec season (Table 1). These baboons live in groups of adult females that are typically relatively closely related to each other, their immature offspring, and one to a few unrelated adult males[54]. Vervets live in similar groups, though often with a greater proportion of adult males per troop[55]. Both species are considered opportunistic and eclectic omnivores[55,56]. These primates typically forage on the ground and in trees during the day, and retire to sleep in trees or on cliffs at night[55] Sample sites were based on known sleeping sites of the primate groups being studied, and were all in the southeastern quadrant of MRC or in nearby Naibor.

### Primate sampling

We obtained fecal samples, group composition data, and movement data for five groups of baboons (six including the supergroup – see above) and six groups of vervets variably over three seasons in the course of one calendar year: January, April-May, and November-December. At least one individual in most of these groups was fitted with a GPS collar for other, unrelated, research studies, and we were thus able to track the locations of these groups daily and know group compositions. We aimed to resample each of these original groups in each of the three seasons, but were sometimes unable to do so due to troop migrations off-site (Table 1). We added two additional groups of baboons in the Naibor area to the sample set in the final season, but were only able to estimate group numbers due to lack of tracking devices in these groups.

We tracked groups of baboons and vervets nearly every evening of each sampling period. The following morning, we noninvasively collected only fresh fecal samples from underneath a group’s sleeping trees or from its sleeping cliffs. Fecal samples were collected between 8:00 and 11:00AM to control as much as possible for temporal parasite shedding variation[57]; as most groups left their sleeping sites between 7:00 and 11:00AM, and many group members defecated prior to leaving (pers. obs.), we are reasonably confident that most samples were freshly shed within this morning window. The sampling sites encompassed seven areas along the Ewaso Nyiro River along the eastern edge of the MRC, two cliff sites (one approximately 1km north of the river, and one approximately 5km west of the river), two sites in human camps (one at the central MRC buildings and one at the Mpala Ranch buildings), one site near a cattle dip in Naibor, and one site in a subsistence garden in Naibor.

Samples were collected into fresh plastic zipper bags, labeled, and put into portable coolers (held at 4C) for 1-3 hours until returning to the laboratory, when samples were placed in a refrigerator at 4C. Parasites were counted from fecal samples starting immediately upon arrival at the laboratory. The majority of samples were clearly from adult animals, based on relative uniformity of size and mass. We labeled significantly smaller (less than 50% typical size in all dimensions) fresh fecal samples as likely being from juveniles.

We aimed to collect samples from at least 50% of group members each season. In nearly all cases, we collected samples from a group all on one morning to best avoid duplicates; these primates tend to defecate just once prior to leaving sleeping sites as a group (pers. obs.), ensuring that each fresh sample in a morning represented just one individual. We collected a total of 318 fecal samples from MRC baboons, a sampling depth of 34.7%, 48.5%, and 48.3% of the baboon population living in this area of MRC during each season respectively. We collected 48 samples from the two groups in Naibor in the final sampling season, 45% and 40% of each group based on rough estimates of group size. We collected a total of 327 fecal samples from vervet monkeys, a sampling depth of 50.0%, 70.1%, and 67.3% for each season respectively based on known group numbers (Table 1).

In almost all cases, only a portion of each troop was sampled during each season. In addition, in nearly all cases, only a portion of each sampling set for each troop was sequenced for microbiome analysis. Given these percentages, there was only a 2%-6% chance of resampling the same individual in two seasons, and a 0.6% chance of sampling the same individual three times across the population. In addition, 5 of the groups were sampled only once, rendering across-season pseudoreplication for those troops impossible (Table 1). Four groups were sampled in only two seasons (another 2 troops were sampled so lightly in one season as to essentially be only tracked over two sampling seasons rather than three). Only 3 troops were sampled enough to capture between 10% and 47% of the individual members for parasite x microbiome analysis over three seasons.

For the 8 troops that were sampled in more than one season, we compared total parasite counts within each group across sampled seasons by Kruskal-Wallis tests or Mann-Whitney tests, and also by Friedman tests and Wilcoxon tests; we did the former set of tests on the assumption that samples were independent across seasons, and the latter set of tests assuming resampling of individuals. We found that total parasite counts were significantly different across seasons for 6 of the resampled troops (p<0.05), and significantly different for another group at p<0.1. These data indicate that the vast majority of troop individuals differed in their parasite infection intensity over sampling seasons. This, combined with the relatively low chance of resampling the same individual for both parasite and microbiome analyses in separate seasons, makes us reasonably confident that any individuals that may have been resampled were likely experiencing differing parasite infections over time, resulting in different microbiome interactions.

### Gastrointestinal parasite species and quantification

The gastrointestinal (GI) parasites examined in these host species included genera within five different groups. Two groups were within the order Rhabditida: a *Strongyloides* species (superfamily Panagrolaimoidea, family Strongyloididae), and seven likely distinct species of strongyle nematodes (suborder Strongylida, primarily within superfamily Strongyloidea and family Strongylidae)[58]. While we were unable to definitively speciate the strongyles in our samples, previous work in baboons in Tanzania, Kenya, and Uganda revealed the presence of *Necator, Oesophagostomum, Trichostrongylus, and Ternidens* genera in fecal samples[59–61]. Other parasite studies have corroborated that these strongyle genera are common in wild baboon species in other areas of Africa[62–67]. While molecular sequencing is now often the method of choice for definitively identifying parasite species, this method is not available to all studies for a variety of reasons (*e*.*g*., availability of equipment and laboratory access, financial limitations); the majority of studies cited here used visual parameters of parasite eggs from fecal samples to identify strongyle parasites to genus level. Similarly, we used visual parameters during parasite counting, guided by these studies and scores of veterinary and public health resources, to identify strongyle parasites to the best of our ability. Based on the following parameters, we hypothesize that our NHPs were host to: *Oesophagostomum* (Strongyle 1) (∼60-75um long x 35-40um wide; later stage of cleavage with more blastomeres with less differentiation compared to Strongyle 2); hookworm, most likely *Necator* based on other baboon studies (Strongyle 2) (∼60-75um long x 35-40um wide; thinner shelled; more differentiated and fewer blastomeres in center of egg compared to Strongyle 1); *Trichostrongylus* (Strongyle 5) (∼80-95um long x 40-50um wide; tapered at one end; sometimes with wrinkled inner membrane); and *Ternidens* (Strongyle 6) (∼80-95um long x 40-60um wide; larger and rounder than Strongyles 1 and 2; often with 4 large morulae filling most of egg). We noted and counted other strongyles that were likely additional genera/species, but were unable to differentiate them further (Strongyles 3, 4, and 7). We identified one tapeworm species (order Cyclophyllidia). We identified one species of *Trichuris* (order Trichocephalida, family Trichuridae) [58]. We also identified two likely distinct species of coccidian parasites (Class Conoidasida, subclass Coccidia). We occasionally observed larvae in our samples that differed from *Strongyloides* larvae that may have burst from eggs during processing; as the fecal samples were collected fresh (within 1-3 hours of defecation), we hypothesize that these may indicate infection with lungworms in some hosts.

We evaluated fecal samples for eggs and oocysts using a modified McMaster technique for fecal egg counts [69]. While the results from this technique can be affected by factors such as season, time of day, host behavior, host genetics, and fecal moisture content[57,70,71], this method has been found to have high sensitivity with lower variation than many other non-invasive parasite sampling methods[72], and is very commonly used to non-invasively quantify parasitism in wildlife and veterinary species lieu of lethal sampling[71,73]. We conducted all parasite counts the same day that samples were collected. We dried a subsample of each fecal sample and adjusted parasite counts to dry weight, as fecal moisture can skew parasite intensity estimates[71]. We controlled for other factors potentially influencing parasite counts by collecting only fresh samples at the same time each morning; accounting for host behavior, diet, and genetics (through group identity, wherein individuals are typically related and share overall dietary and behavioral preferences – see Methods, above); and accounting for season of collection in our analyses. We collected data on parasite infection intensity (as eggs per gram, epg), infection richness, and infection composition.

### Microbial DNA extraction from feces and sequencing

We chose fecal samples for microbial DNA extraction with the aim of representing at least 30% (when possible) of a group’s samples in our sequencing efforts (Table 1). When we needed to reduce the number of samples chosen from a group (*i*.*e*., a group had more samples than were economically feasible to sequence), we chose samples on either extreme of the parasite infection intensity scale (*i*.*e*., those with no or few parasite epg, and those with the highest epg shedding), and those on either extreme of the parasite richness scale (*i*.*e*., those with monospecific infections and those with the most diverse infections; we chose samples from these groups by starting with the outliers on each end of the data extremes and “working back” toward those samples falling within quartiles 1-3; S1 Fig). Parasites are typically overdispersed within a population[74]; they tend to be very abundant in a few individuals in a population, and at much lower levels in the rest of a group, a phenomenon we observed within our NHP family groups (S1 Fig). We were thus able to use this as a natural experiment. While we accounted for helminthic abundance in our phylofactorization models (see below), we were essentially able to compare “very wormy” individuals in a group to “control” (*i*.*e*., those with few to no parasites) animals within the same group. Given that most animals in a stable group are related, this helped us to control for genetic variation and physiological background while focusing as much as possible on the correlations between parasite presence and microbiome composition.

We extracted microbial DNA from fresh fecal samples using Zymo Xpedition Soil/Fecal DNA Mini Prep Kits with Zymo DNA/RNA Shield (Zymo Research, Irvine, CA); fecal samples were kept in DNA/RNA Shield in a refrigerator for no more than four days (96 hours) prior to extraction[75]. We measured DNA yields with Qubit Fluorometric Quantitation (Life Technologies, Grand Island, NY). We did PCR amplification with 16S rRNA primers for the V4 region of *E. coli* (primers F515 and R806) to determine which samples yielded usable DNA. We sent the usable samples to Michigan State’s Research Technology Support Facility (East Lansing, MI), which built 16S libraries using primers for the 16S rRNA V4 region[76] by Illumina MiSeq sequencing of 2×250bp end reads; OTUs were assigned by closed-reference OTU picking with a 97% sequence similarity cutoff and the resulting OTU table rarefied to 8,000 reads per sample by random subsampling. We sequenced a total of 143 baboon samples from MRC (representing 40.0%, 39.5%, and 57.1% of the entire MRC baboon population across each of the three seasons, respectively) and 143 vervet samples (representing 23.9%, 38.8%, and 45.0% of the total vervet population across the three seasons; Table 1). We sequenced 23 samples from the two Naibor sites, representing 17.5% and 30.0% of the two groups in that area.

### Phylofactorization and statistics

We used phylofactorization[20,21] of the microbial community dataset for flexible identification of microbial clades associated with host species and helminth infections. All scripts for these analyses are available upon request. Phylofactorization discovers lineages of microbes that maximize an objective function, such as associations between their relative abundance and helminth presence, controlling for confounds such as species and month. We chose to use phylofactorization out of a prior understanding that some important microbial functions will be shared among close microbial relatives whose evolutionary history is reasonably well summarized by the evolution of housekeeping genes, and that lineages of microbes found to be associated with helminthic colonization of primates in this study can be tested for their functional associations in other studies by their common phylogenetic scaffold.

Here, we use the same Greengenes tree as our phylogenetic scaffold as was used in the original phylofactorization analysis of the American Gut Project (AGP)[20]. OTUs with fewer than 3 counts across all samples were excluded from this analysis. The AGP analysis with phylofactorization found that a large lineage, made up almost entirely of *Prevotella* OTUs, was associated with inflammation. In our current study, we find that a large lineage, made up almost entirely of *Prevotella* OTUs based on the same Greengenes taxonomy, is associated with helminth infections.

To identify major axes of variation in the microbial community attributable to species differences, we performed phylofactorization using the R package phylofactor, downloaded from https://github.com/reptalex/phylofactor. The datasets analyzed during the current study are available from the corresponding author on reasonable request and will be made available in on online repository after completion of further analyses by our team.

We first performed phylofactorization by maximizing the explained variance of phylogenetic component scores, *ye*, for each edge e, by species through the model *ye*∼species.

To identify clades of microbes with the most predictable responses to helminths, and put helminths on a comparable scale, we converted helminth abundances into presence/absence and performed phylofactorization using the formula

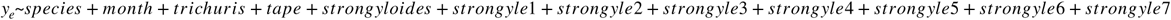

and maximized the deviance from the “worm” model containing species, month, *Trichuris*, Strongyle 2, tapeworms (tape), and all other strongyles (totalstrongyle) relative to the model containing only “species” and “month” (see S2 Fig). We limited our presentation to clades of microbes – as opposed to single OTUs – as hypothesized functional groups of species that can be compared in other model systems using different OTUs within the same lineage (*e*.*g*. the murine model for inflammation may utilize strains of *Prevotella* not found in our specific survey of primates, but can be determined to fall in the same crown group).

To control for multiple hypothesis tests, we computed P-values from an F-test of the deviance of models with vs. without the “worm” terms and used Holm’s conservative sequentially-rejective procedure[77], as discussed in the ecological monograph on phylofactorization[20], to correct P-values. We present only large clades whose corrected P-values fall under a 5% family-wise error rate threshold for Holm’s sequential Bonferroni procedure.

Holm’s sequentially rejective procedure has been shown to be a conservative procedure for phylofactorization as its assumptions of independence between neighboring edges overstates the effective number of multiple comparisons. By considering only factors that fall under a 5% family-wise error rate (FWER) with Holm’s procedure, we are reporting only on a set of lineages with a <5% chance of there being a single false-negative in our set of lineages. We do not report raw P-values of lineages because: (1) these P-values from F-tests of the deviance at every iteration are the lowest of the set of P-values for every edge in the tree (hence reporting a q-value or the minimum FWER at which the lineage would be included in a set of positive results might be more appropriate); and, more importantly, (2) due to the dependence between edges in the tree, the assumptions of independence in FWER and FDR calculations do not hold. Consequently, these P-values, q-values, minimum FWERs at which a result would be considered significant, and other multiple comparison and multiple hypothesis-testing procedures are not properly calibrated for phylofactorization at this time (similarly, these methods are also not properly calibrated for factor analysis, including for PCA). Consequently, we limit our reporting only to a set of lineages with <5% FWER to give a familiar, conservative, and correct “P<0.05” intuition that applies to all the phylogenetic factors we report.

## Supporting information

Supplemental Material - Microbial Sequence Database

Supplemental Material - Parasite Database

## Acknowledgements

We thank the Mpala Research Centre in Nanyuki, Kenya for allowing the field portion of this work to be conducted on its premises. We thank L. Bidner and L. Isbell for primate group background information and tracking assistance; B.K. Sitienei for assistance with sample collection and initial sample processing; B. vonHoldt for PCR amplification expertise and assistance; and D. Rubenstein for field vehicle use.

## Supplemental information

**Supplemental Figure 1.**
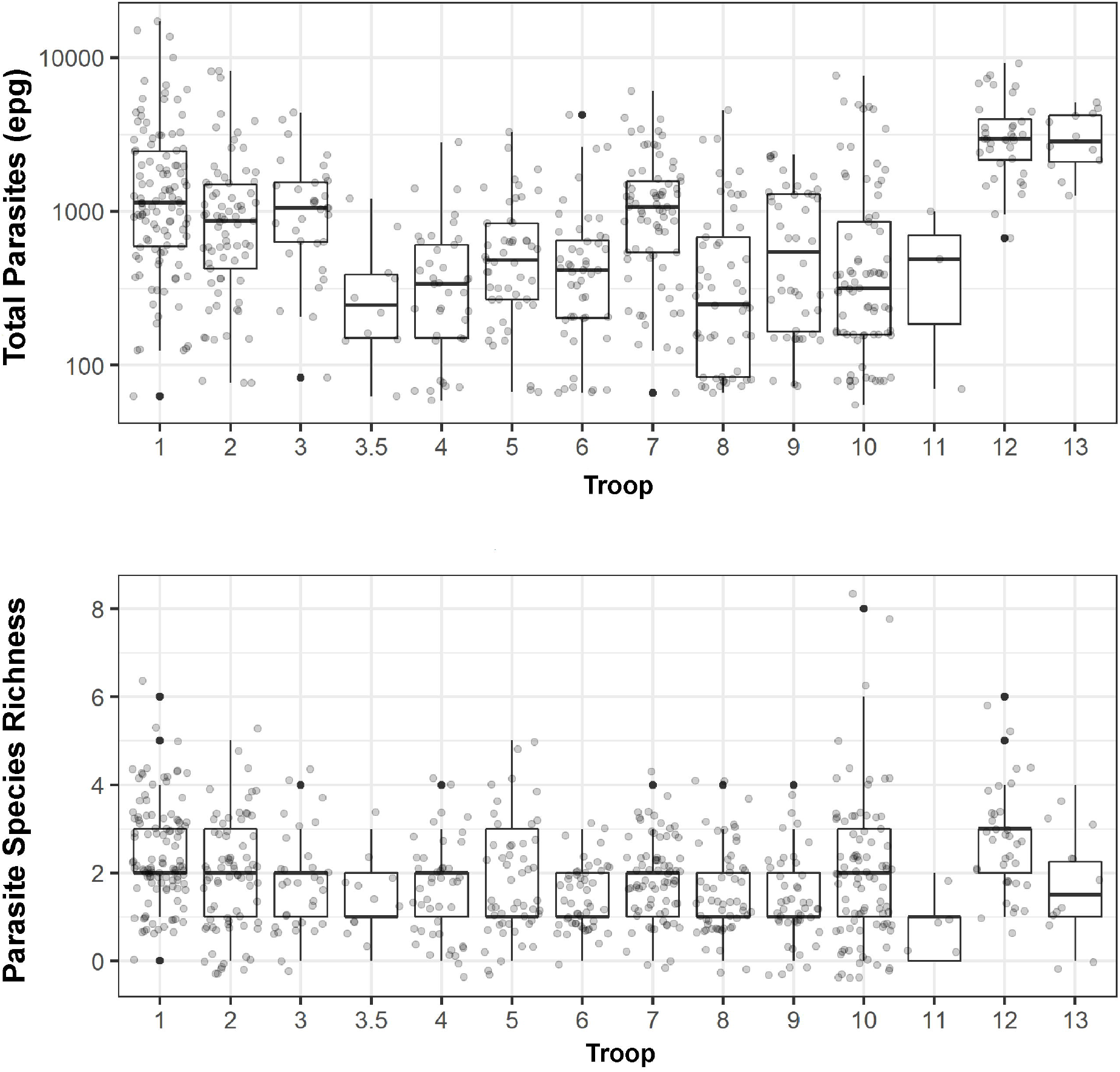
Total parasite counts and parasite genera/species richness by troop identity. These boxplots reveal that most of these parasites are highly aggregated in their hosts (indicated by outliers), with the majority of parasites existing in the minority of hosts in each troop. The majority of individuals have intermediate parasite richness (maximum richness is 13 genera/species). Troop identities are as follows: 1 Cliffs baboons; 2 Meg’s baboons; 3 Shorttail’s baboons; 4 Limpy’s baboons; 5 Fig vervets; 6 Ranch vervets; 7 Hippo Pool vervets; 8 Bridge vervets; 9 Small baboons; 10 Kudu vervets; 11 MRC vervets; 12 Naibor Cattle Dip baboons; 13 Naibor Garden Site baboons.

**Supplemental Figure 2.**
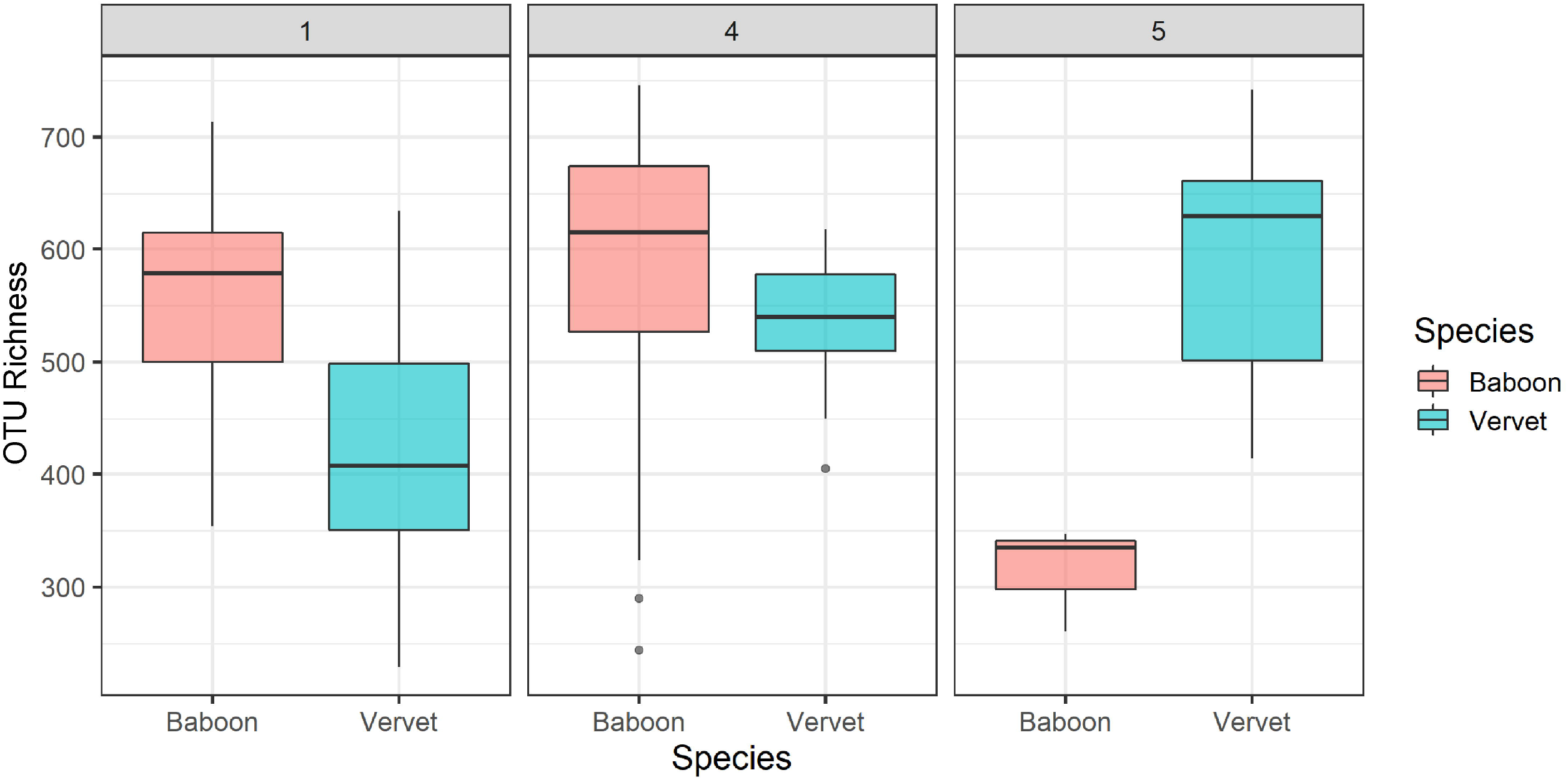
OTU richness (number of OTUs) across host species and month. Due to significant differences of microbiome composition across species {Baboon,Vervet Monkey} and months {1,4,5}, we focus phylofactorization on worm-associated lineages when controlling for categorical factors of species & season.

